# A benchmark study of data normalisation methods for PTR-TOF-MS exhaled breath metabolomics

**DOI:** 10.1101/2023.06.22.546053

**Authors:** Camille Roquencourt, Elodie Lamy, Emmanuelle Bardin, Philippe Deviller, Stanislas Grassin-Delyle

## Abstract

**Background:** Volatilomics is the branch of metabolomics dedicated to the analysis of volatile organic compounds (VOCs) in exhaled breath for medical diagnostic or therapeutic monitoring purposes. Real-time mass spectrometry technologies such as proton transfer reaction mass spectrometry (PTR-MS) are commonly used, and data normalisation is an important step to discard unwanted variation from non-biological sources, as batch effects and loss of sensitivity over time may be observed. As normalisation methods for real-time breath analysis have been poorly investigated, we aimed to benchmark known metabolomic data normalisation methods and apply them to PTR-MS data analysis.

**Methods:** We compared seven normalisation methods, five statistically based and two using multiple standard metabolites, on two datasets from clinical trials for COVID-19 diagnosis in patients from the emergency department or intensive care unit. We evaluated different means of feature selection to select the standard metabolites, as well as the use of multiple repeat measurements of ambient air to train the normalisation methods.

**Results:** We show that the normalisation tools can correct for time-dependent drift. The methods that provided the best corrections for both cohorts were Probabilistic Quotient Normalisation and Normalisation using Optimal Selection of Multiple Internal Standards. Normalisation also improved the diagnostic performance of the machine learning models, significantly increasing sensitivity, specificity and area under the ROC curve for the diagnosis of COVID-19.

**Conclusions:** Our results highlight the importance of adding an appropriate normalisation step during the processing of PTR-MS data, which allows significant improvements in the predictive performance of statistical models.

## 1. Introduction

Medical metabolomics is the science of small molecules in the human body and the data generated during mass spectrometry metabolomic analysis consists of the abundances of metabolites measured in biological samples under different conditions (factors of interest) [1]. Depending on the technique used to generate the data, the data must be pre-processed with the aim of subsequent statistical analysis to uncover metabolites with different abundances between the biological factors of interest [2]. However, unwanted variations in metabolite abundances can occur during data collection, either due to instrumental variation (drift in mass accuracy, sample degradation, loss of sensitivity over time…) or inter-instrumental variations (different laboratory environments, different analytical platforms…). Data normalisation therefore aims to reduce the variation from non-biological sources, while preserving the biological variation of interest unaffected [3–5]. Several normalisation methods for metabolomic data have been described and can be divided into three categories [6]: (i) statistical methods based on the complete data set, applying an individual scaling factor to each sample [7]; (ii) internal standard (IS)-based methods, where the IS are (exogenous) metabolites whose expression does not vary with the biological factors of interest. They are usually added to the sample in a known quantity prior to analysis so that they can reflect all sources of unwanted variation independent of sample collection and storage; and (iii) normalisation using quality control (QC) samples, which are a combination of equal volumes of each sample with an overall composition that is very comparable to the study samples.

A branch of metabolomics is volatilomics, the analysis of volatile organic compounds (VOCs) in exhaled breath, which is of great interest for the development of non-invasive diagnostic methods [8]. Several real-time technologies are available for breath analysis, including proton transfer reaction time-of-flight mass spectrometry (PTR-TOF-MS) [9,10], which detects and measures the mass-to-charge ratio (*m*/*z*) of VOCs present in a sample with high sensitivity and specificity, provided that they can be ionised. In brief, ionisation is the first step based on proton transfer from a reagent ion, most commonly hydronium ions generated from a water source, before the mass of the ions can be measured by their time of flight from the transfer chamber exit to the detector. An advantage of PTR is the conversion of ion intensities into “absolute” quantities, which is achieved by normalising the ion intensities by the intensity of the reagent ion (i.e. *H*_3_*O*^+^), the reaction rate coefficient between the VOC and the reagent ion, and the residence time of the primary ions in the drift tube [11] [12] [13]. The final normalised concentration is then obtained by further division by the density of the air in the reaction chamber, expressed in parts per billion (ppb). This normalisation method can be considered as a single standard normalisation as the reagent ion is continuously generated and analysed during the acquisition. However, normalisation with a single molecular component is very sensitive to its own variation and is not always sufficient to remove all unwanted variation. As is common in mass spectrometry analysis, a time-dependent loss of intensity can occur over long periods of time. Although some of this analytical drift is related to the response of the microchannel plate (MCP) detector and some can be corrected by adjusting the voltages, which in turn reduces the life of the detector, the degradation in performance can still occur and needs to be corrected.

The aim of this study was first to evaluate signal intensity drift in PTR-TOF-MS analysis and then to benchmark available metabolomic normalisation methods to propose the most suitable for exhaled breath analysis by PTR-TOF-MS. Quality control based methods were excluded as not suitable for real-time breath analysis.

## 2. Methods

### 2.1 Normalisation methods

We first identified then benchmarked the normalisation methods commonly reported for classical metabolomic datasets from liquid or gas chromatography MS analysis of biological samples such as plasma, urine or tissues [14–17].

We denote *X* a *n* × *p* matrix of signal intensities in count per second (cps) units, with *n* samples in rows and *p* VOCs in columns. Missing data imputation was performed as a prerequisite for all normalisation methods (described in 2.3). We denote 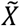 the normalized matrix, *X*_*i*_ the *i*^*th*^row (sample) and *X*_*j*_ the *j*^*th*^ column (feature).

The normalisation methods are the following:

- **Total sum of the signal (TSS)**, consisting in dividing the intensity of each feature by the sum (or the sum of square) of the intensities of all the features in the sample. This makes the hypothesis that the total amount of VOCs detected is identical in all samples. The main limit of this method is the greater influence of the most intense VOCs on the normalisation.

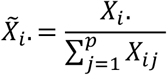
- **MS - total useful signal (MSTUS)**, uses as a denominator the intensity of VOCs that are present in given proportion of all samples (e.g. > 95%) and only in the background air (not in breath phases, to avoid to discard the biological variation of interest). The inter-sample variance of these features is supposed to be low, since it represents VOCs measured during ambient air analysis.

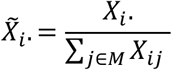

with *M* = {*j*, s.t *X* _*j*_ have missing value <5% of the samples and is not selected as exhaled breath VOC}.

- **Median normalisation** divides the intensity of each feature by the median of the total signal [2]. In contrast to TSS, it is not sensitive to extreme values, especially in situations where several saturated features may be associated with some of the factors of interest.

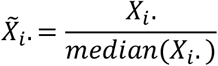
- **Quantile median normalisation (QM)** aims at achieving the same distribution of feature intensities across all samples [18]. The features are first ordered by intensities for each sample *i*: (*X*_*i*_[1], … *X*_*i*[*p*]_). Then the mean of each column is computed, which corresponds to the mean of the *j* ^*th*^ more intense VOC across sample: 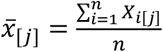, and finally, the original order of the assigned mean values is restored to determine the normalized intensity 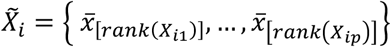.
- **Probabilistic quotient normalisation (PQN)** scales by analysing the distribution of the quotients of the intensities in each sample out of the intensity in a reference sample [19]. It means that a reference and unchanging sample has to be selected and used to rescale the remaining samples by the median of the quotient:

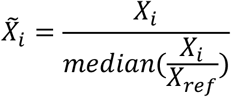

The reference sample is usually the median of quality control samples, when available. For real-time analysis, we chose a reference sample arbitrarily.

- **Normalisation using optimal selection of multiple internal standards (NOMIS)** finds optimal normalisation factor to remove unwanted variation using multiple compounds as internal standards [20]. This method selects the best combination of internal standards using multiple linear regression. The starting hypothesis is that the VOC intensities are a multiplicative model of unwanted variation, represented through internal standards, as there are unaffected by the biologic state of interest, allowing to assume that any variation of these metabolites is only arising from unwanted interferences:

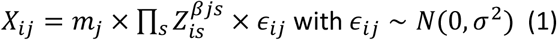

where *z* is an independent matrix of *X*, containing the intensities of *S* standards, and *m*_*j*_ is the true intensity value of the VOC *j*, independent of sample *i*. This assumption for *m*_*j*_ means that independent measures from one biological specimen (e.g. under repeatability conditions) have to be performed for training the normalisation coefficients *β*, which value can then be applied for the normalization of new datasets, even for different biological specimen. For the application to real-time breath analysis, we propose to use VOCs from several measurement of the same background air over days. A linear regression is then applied to the log-transformed model (1) after mean-centring the matrix Z of log internal standard intensities:

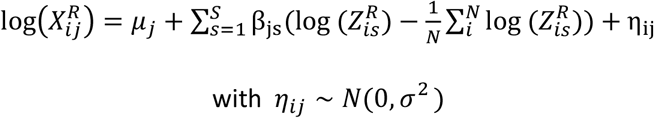

where *X*^*R*^, *Z*^*R*^ are the matrix of room air intensities, analysed the same day than each exhaled breath sample. The *β*_*js*_ coefficient then represents the impact factor of the internal standard *s* on the VOC *j*. When the parameters have been trained on ambient room air, it can be applied to the whole clinical cohort *X*, but the same internal standards must be also present in the patient samples. The normalised matrix is finally obtained by dividing the product of standard intensities, weighted by the trained coefficients 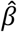:

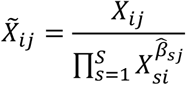

However, this method requires standards and independent repeated measurements, which must be considered in the study design, prior to sample analysis.

- **Remove unwanted variation (RUV)** [7]. As NOMIS, this method requires a set of VOC standards, which reflect the unwanted variation and are independent from the biological state of interest. First, a factorial analysis of log-transform internal standard intensities is performed:

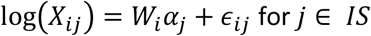

where *W*_*i*_ and *α*_*j*_ are the results of the factorial analysis with *K* factors. Then, the variation estimated by the factor analysis on the IS, 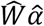 is subtracted from the intensity matrix *X*:

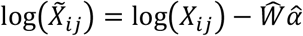

An improvement of this method has been developed, called RUV-random, by adding an iterative estimation of the factorial analysis *Wα* [21]. It makes the hypothesis that:

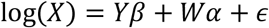

with *Y* containing the biological factor of interest and *W* the unwanted variation. The initialisation starts with the same factor analysis than the RUV method, then the two following steps are iterated: (i) performs a k-nearest neighbours (KNN) algorithm on the normalized matrix 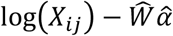 and then (ii) estimates 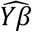 to obtain a refined estimate for α given the estimates for W, using 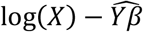, until convergence of α.

### 2.2 Datasets

The datasets from two previous studies were used to benchmark the normalisation methods. The first dataset consists of exhaled breath analysis from participants undergoing COVID-19 screening in the Emergency Department (ED), with data collected over a 12-month period between March 2021 and June 2022 [22]. The second dataset includes analysis of exhaled breath from intubated, ventilated patients with COVID-19-related acute respiratory distress syndrome in the intensive care unit (ICU) between March and June 2020 [23]. The two study protocols were registered (VOC-COVID-Diag: EudraCT 2020-A02682-37; RECORDS trial: EudraCT 2020-000296-21) and were approved by an independent ethics committee. Written, informed consent was obtained from all the participants. Background air from the PTR-TOF-MS room was analysed prior to each sample for the ED cohort and was used to train the normalisation coefficient of the NOMIS method, while medical air was used for the same purpose in the ICU cohort.

### 2.3 Data pre-processing

Mass spectrometry data were processed using the ptairMS R package [24]. Breath was differentiated from background or medical air using acetone (*m*/*z* 59.049) and *CO*_2_ (*m*/*z* 44.997) as tracer VOCs, respectively. Mass calibration was performed every minute using peaks at *m*/*z* 21.022, *m*/*z* 60.053, *m*/*z* 203.943 and *m*/*z* 330.850. After an alignment step, features present in at least 60% of one group and with a signal intensity significantly different between room air and exhaled breath in at least 30% of the samples were retained. Missing data were then imputed from the raw data; isotopes and saturated ions (*m*/*z* 37.028 and 59.049) were removed as well as outlier samples, defined as those with a Z-score > 5 for at least 10% of the features. Finally, the data were log-transformed.

### 2.4 Choice of standard VOCs

As described above, the NOMIS and RUV methods require a set of internal standards. In classical mass spectrometry analysis, internal standards are usually isotopically labelled metabolites added to the sample prior to analysis. Their concentrations are in no way correlated with the biological factor of interest, allowing the unwanted variation to be observed. However, adding internal standards to the sample was not suitable for our real-time analysis platform, so we decided to use the following VOCs, which result from the ionisation process or are naturally present in ambient air, as internal standards:

- **Water-derived ions**: The generation of hydronium *H*_3_*O*^+^ ions from water for the proton transfer reaction also produces the water clusters (*H*_2_*O*)_2_ + *H*^+^ and (*H*_2_*O*)_3_+ *H*^+^, which were studied as internal standards. As the detector is saturated when analysing VOCs containing ^16^O, the most abundant isotope of oxygen, we used stable ^18^O isotopes for these VOCs, corresponding to *m*/*z* 21.022, 39.03 and 55.04 respectively.
- **VOC correlated to the reagent ion**: As the reagent ion can already be used as a single standard for PTR-TOF data normalisation, we investigated the use of multiple standards by selecting all VOCs whose intensities across the sample correlated with the reagent ion according to a Spearman correlation test (p-value <0.001).
- **Useful signal**: Inspired by the MSTUS method, VOCs are selected as internal standards if they are present in more than 95% of the samples and if they are not selected as VOCs from the exhaled breath phase.

### 2.5. Metrics

Several metrics were calculated to benchmark the normalisation methods and to evaluate the effect of removing unwanted variation on the performance of predicting the biological factor of interest:

- Linear regression between geometric mean of intensities in each sample and acquisition date to assess time-intensity relationship and signal correction. We used the geometric mean (the exponential of the log mean) because it is less sensitive to the highest values in a data set than the arithmetic mean.
- Predictive metrics of the scientific question relevant to the underlying study objective: sensitivity, specificity, and area under the ROC cure (AUC) for the diagnosis of COVID-19, using a Random Forest model with stratified 10-fold cross validation repeated 2 times and grid search on the number of variables randomly sampled as candidates at each split.

## 3. Results

Data were available for 139 participants in the emergency department cohort (41 COVID-19 positive and 98 COVID-19 negative) and 29 patients in the intensive care unit cohort (19 COVID-19 positive and 10 COVID-19 negative). In the ED cohort, a total of 347 features were detected during data pre-processing, including 67 features with a higher expression level in breath samples than in background. For the ICU cohort, the corresponding number of features was 195 and 105, respectively. Only features from exhaled breath were used for statistical analysis and categorisation of patients into COVID-19 positive or negative groups, but all VOCs (from breath and background) were used in the normalisation algorithms. A plot of the raw data in each cohort is shown in Figure 1.

**Figure 1:**
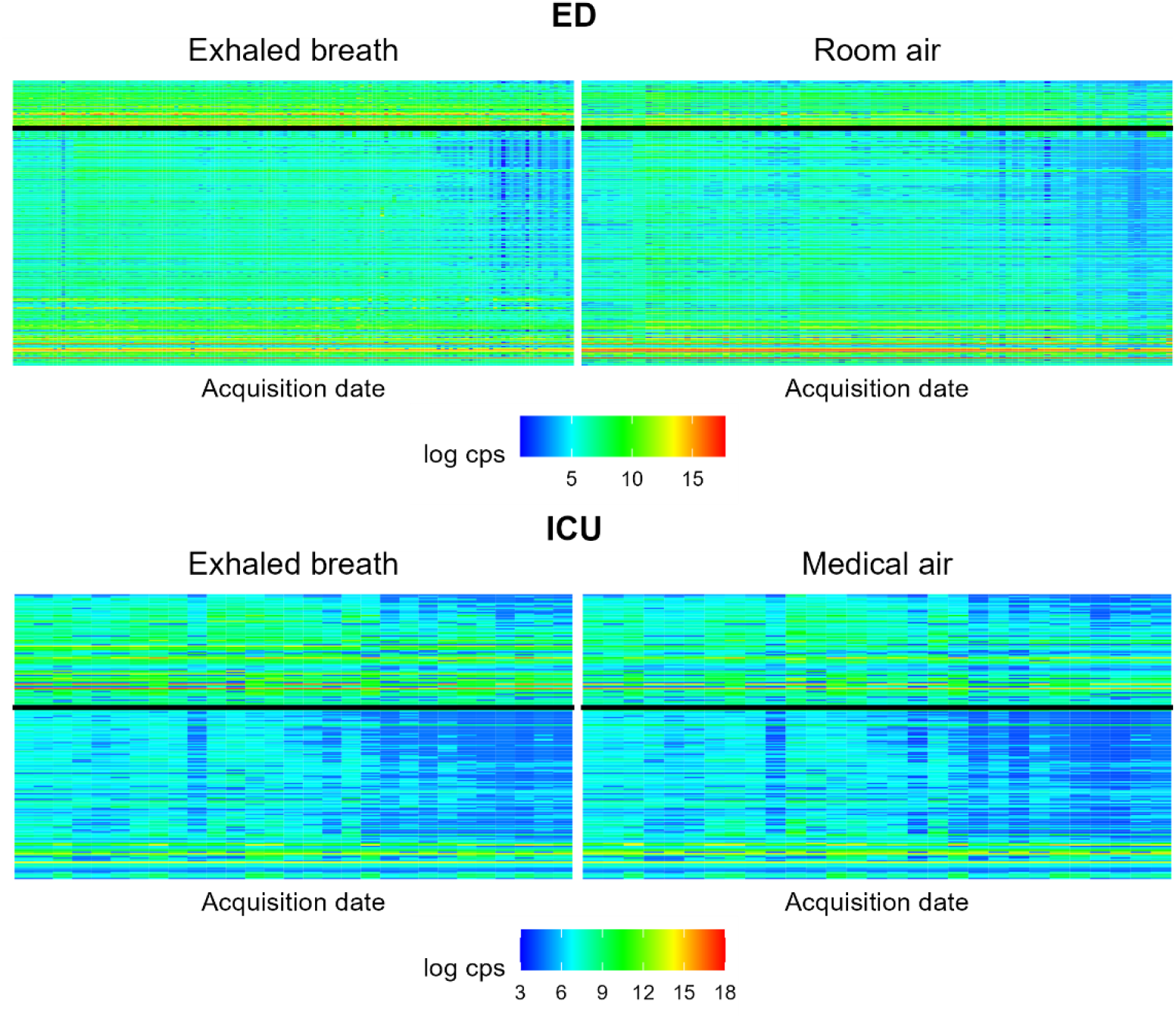
Graph of peak tables for the emergency and ICU cohorts. The left panel represents the patient data and the right panel represents the background air (room air or medical air) used to train the NOMIS normalisation coefficients. VOCs above the horizontal black line correspond to VOCs with a significantly higher expression level in breath than in background air. Data were expressed in counts per second (cps) and log transformed.

Drift in signal intensity using uncorrected raw data was first assessed using linear regression between the geometric mean of intensities from each sample and the date of acquisition. The results in Figure 2 show a linear decrease in signal intensity over time, with the slope of the regression line being -0.068 cps/month (*p* < 2.10^−16^) for the ED cohort and -0.226 cps/month (*p* = 2.10^−5^) for the ICU cohort.

**Figure 2:**
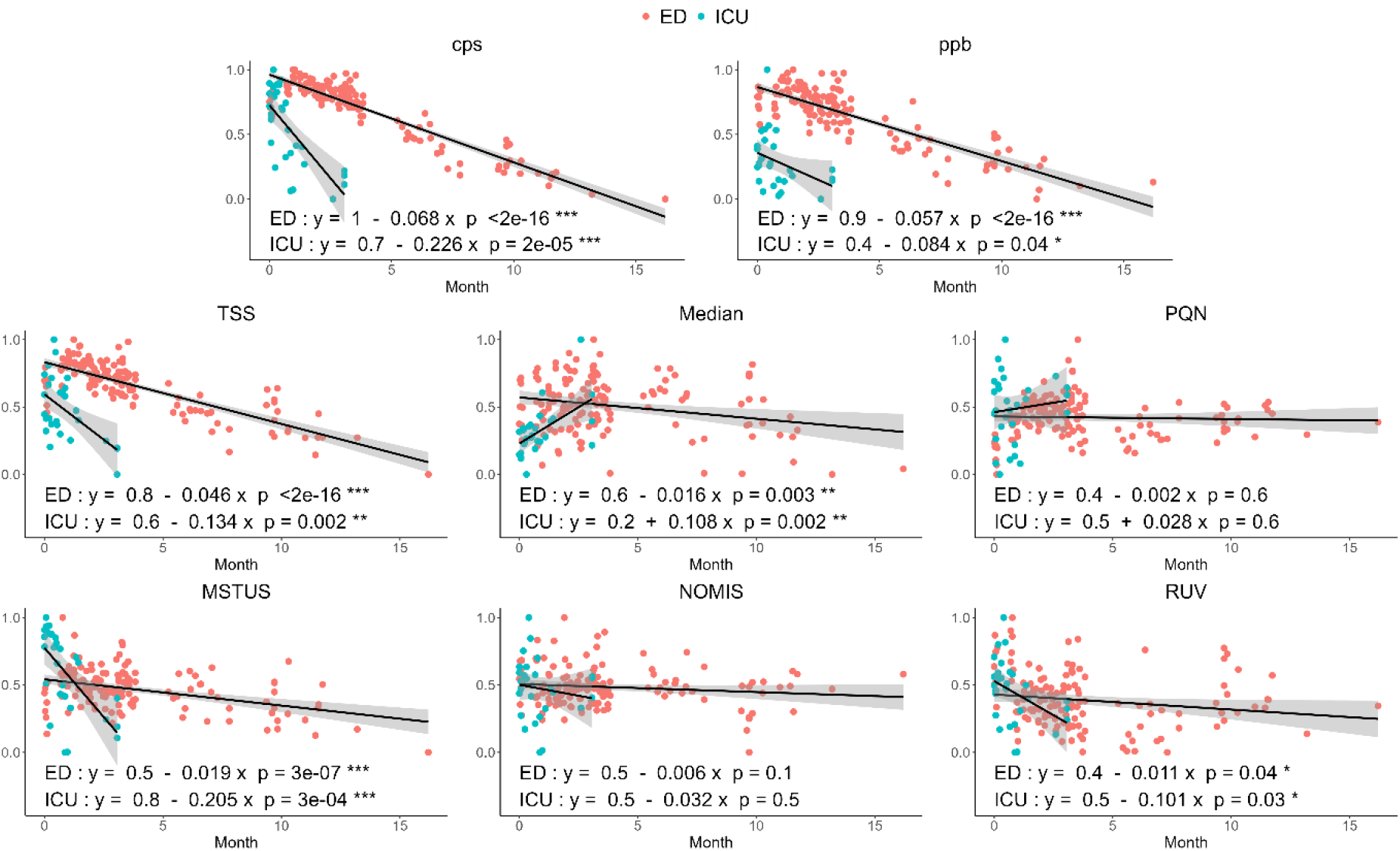
Geometric mean intensity value of each sample normalised between 0 and 1 as a function of acquisition date for the ED and ICU cohorts. Each panel represents the raw data (cps) or each normalisation method. The black line represents the linear regression with the 95% confidence interval in grey. The QM method is not shown as all samples have the same feature distribution.

We then compared the seven normalisation methods and the results are shown in Figure 2. Only the best method (of the 3 methods described in 2.4) is shown for NOMIS and RUV, using AUC as the endpoint, and detailed results are shown in Supplementary Table 1. The results of the QM method are not shown because all samples have the same distribution of feature intensities and therefore the same mean.

The ppb normalisation method slightly attenuated the slope of the signal drift, which nevertheless remained significantly different from 0 (0.057 cps/month (*p* < 2.10^−16^) for the ED cohort and -0.084 cps/month (*p* = 0.04) for the ICU cohort). Detailed results for the 6 other methods are shown in Table 1. Among these, the methods providing the best corrections for both the ED and ICU cohorts were PQN and NOMIS, with slopes between -0.032 and 0.028 cps/month, not significantly different from 0. Regarding the selection of the best standard VOCs for the NOMIS and RUV methods, the use of the three water ions gave the best results for both methods in the two cohorts and for the RUV method in the ED cohort. In this last cohort and for the NOMIS method, the correlation method was best with 29 standards selected and the estimated coefficients for each standard selected are shown in Supplementary Figure 1. For both cohorts, the RUV random algorithm with KNN algorithm during iterations gives a better AUC than classical RUV.

**Table 1:**
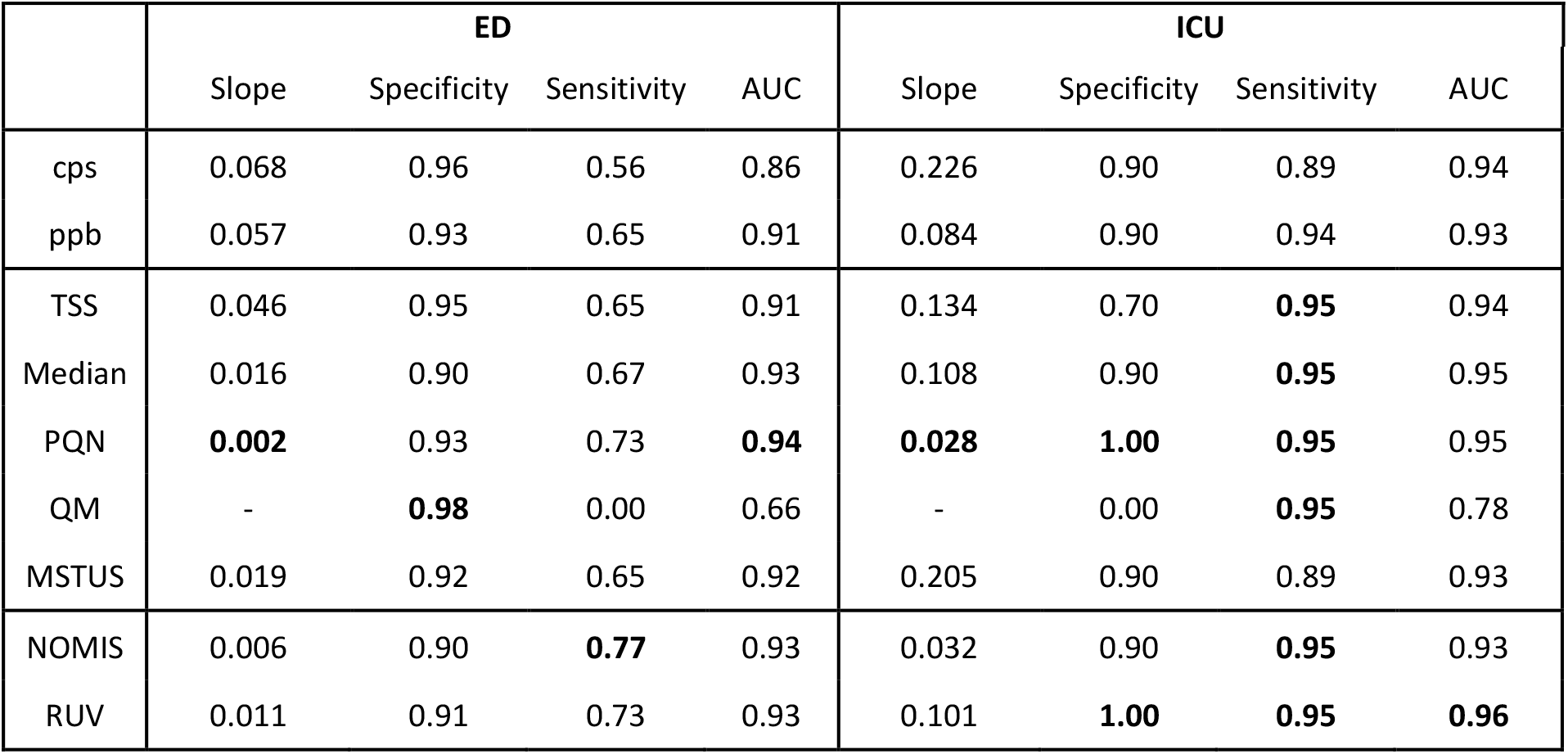
Benchmark of normalisation methods. The different metrics for the different normalisation methods are shown for the ED and ICU datasets: slope of the linear regression between geometric mean intensity value and acquisition date of each sample; specificity, sensitivity and AUC of the random forest model with 10-fold cross validation repeated twice with COVID-19 status as predictor variable. The slope for the QM method is not shown as all samples have the same feature distribution

We then focused on the predictive performance of the models used to classify patients according to their COVID-19 status, after data normalisation with each of the different methods. The results of sensitivity, specificity and area under the ROC curve are detailed in Table 1 and the ROC curves are shown in Figure 3.

**Figure 3:**
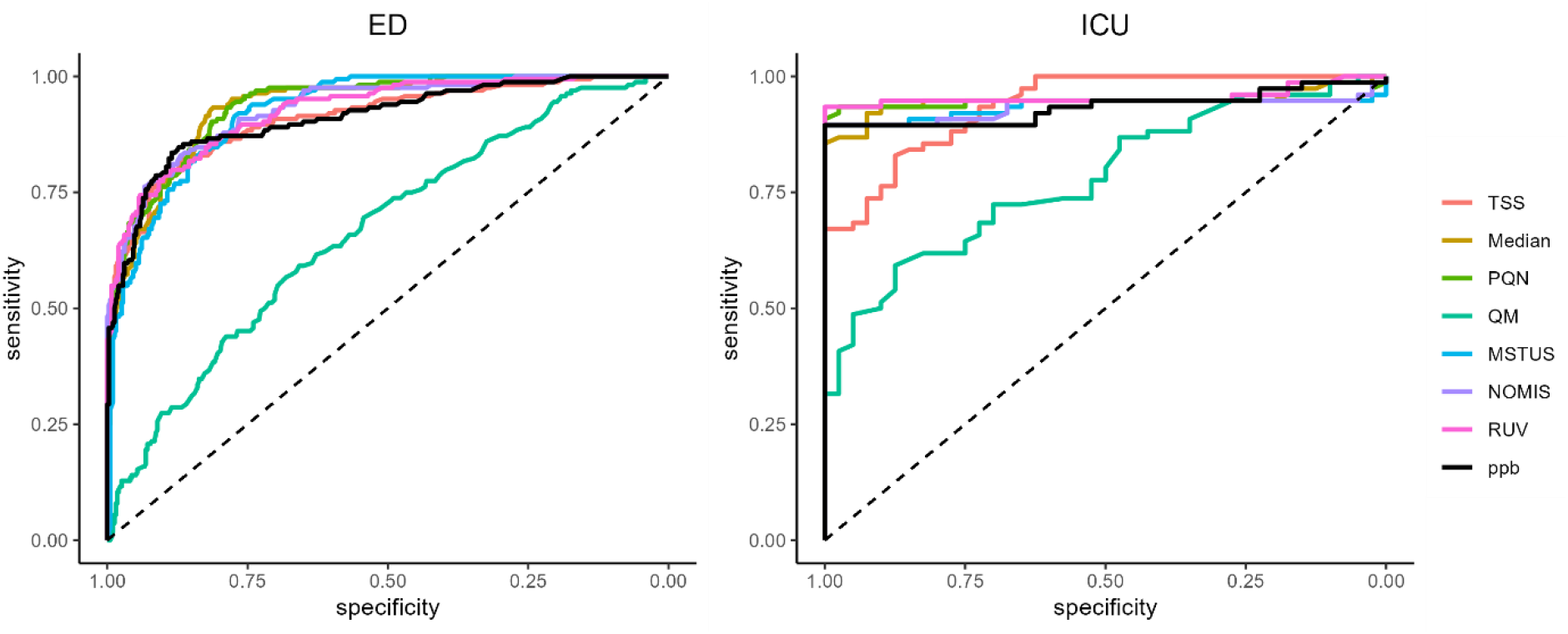
Receiver Operating Characteristic (ROC) curve of each Random Forest model trained with the different normalisation methods using 10-fold CV repeated 2 times.

For the ED cohort, sensitivity increased from 0.56 without normalisation to 0.77 with the NOMIS method, while specificity ranged from 0.90 (median and NOMIS) and AUC from 0.66 (QM) to 0.94 (PQN). The method with the best sensitivity with a suboptimal AUC (0.93) was the NOMIS method with 29 standard VOCs. For the ICU cohort, sensitivity was 0.89 without normalisation and up to 0.95 with 6 of the normalisation methods, including NOMIS, PQN and RUV. Specificity ranged from 0.70 to 1.00 for all methods except QM (0.00), with AUC ranging from 0.78 (QM) to 0.96 (RUV). RUV and PQN were the best performing methods.

## 4. Discussion

Our results show that the time-dependent drift in PTR-MS signal intensity observed during breath analysis of clinical cohorts can be corrected using normalisation algorithms, which in turn improves the predictive performance of models used for diagnostic purposes.

First, the slope of the regression line describing the decrease in signal intensity was greater in the ICU cohort than in the ED cohort. This suggests that the drift is not only related to the time-dependent decrease in detector sensitivity (detector aging), but also to patient- and sampling-related factors such as differences in breath sampling methodology, differences in the quality of the background air (room air vs. medical air) and patient characteristics (awake vs. intubated, ventilated patients; severity of illness…). However, signal correction with the different algorithms was possible and equally effective in both cohorts.

We then benchmarked several normalisation methods. Of all the algorithms tested, those based on statistics (ppb, TSS, median, PQN, QM and MTUS) are easy to calculate and apply to new data sets. The other two (NOMIS and RUV) rely on the use of several standard VOCs, which have to be selected according to specific criteria. These methods may be sensitive to the selected standards, as the three standard selection methods may give slightly different results (Supplementary Table 1). Two of these methods do not require chemical interpretation of the standards and are based only on correlation with the primary ion or reproducibility in ambient air. However, the use of unknown VOC standards should be used with caution as the data are normalised so that the standard concentrations are constant over time, which cannot always be verified as these VOCs may also originate from environmental contaminants [25,26]. We also proposed to train the multi-linear regression of the NOMIS algorithm by using measurements of the same indoor air at the time of sampling for each patient. However, although this method estimates the best linear combination of VOC standards to explain the variation of all other compounds, care should be taken if the measurements used for training are not obtained in the same condition. In these circumstances, the use of the three water ions as standards (m/z 21.02, 37.02 and 55.04) should be more reliable as they are derived from the water used as the reagent for VOC ionisation.

Finally, we observed that the predictive performance of the diagnostic models also improved after the normalisation procedure with the different algorithms. The PQN and NOMIS methods, which already provided the best signal correction, also improved the predictive performance. However, the use of multiple standards in the NOMIS method did not provide much improvement over the statistical PQN method. The improvement in prediction performance was moderate for the median and RUV methods, but superior to ppb normalisation for both cohorts, while the TSS and MSTUS methods did not perform well. This may be explained by the high abundance of the water ion, which contributes a large proportion of the total signal in exhaled breath analysis and therefore correlates with the sum of all VOC intensities used in the TSS method. The QM method was not suitable for data normalisation.

The strengths of our study are the use of two different patient cohorts from COVID-19 clinical trials conducted in two different hospitals, and the comparison of several normalisation methods with different approaches (statistical and standards-based). The main limitation is the need for validation in external independent cohorts.

## 5. Conclusion

In conclusion, this manuscript reports the first benchmark study of metabolomic data normalisation methods applied to exhaled breath PTR-TOF-MS analysis. Our results highlight the importance of adding a normalisation step during the processing of PTR-TOF-MS data, which allows for significant improvements in the predictive performance of subsequent statistical models used for diagnostic purposes.

## ACKNOWLEDGEMENTS

The authors would like to thank all the staff members of the emergency department and clinical research team involved in the study of Foch Hospital and intensive care unit at Raymond Poincare Hospital for their collaboration.

## RESEARCH FUNDING

This work was supported by Agence Nationale de la Recherche (COVINOse, ANR-21-CO12-0004 and RHU4 RECORDS, Programme d’Investissements d’Avenir, ANR-18-RHUS-0004); Région Île de France (VolatolHom, SESAME 2016 and MeLoMane, DIM 1HEALTH 2019) and Fondation Foch.

## ETHICAL STATEMENT

The two study protocols were registered (VOC-COVID-Diag: EudraCT 2020-A02682-37; RECORDS trial: EudraCT 2020-000296-21) and approved by an independent ethics committee. Written, informed consent was obtained from all the participants.

## SUPPLEMENTARY MATERIAL

**Supplementary Table 1:** Results obtained with the NOMIS and RUV methods using the different standard selection methods.

|  |  | ED |  |  | ICU |  |  |
| --- | --- | --- | --- | --- | --- | --- | --- |
|  |  | Specificity | Sensitivity | AUC | Specificity | Sensitivity | AUC |
| NOMIS | US | 0.95 | 0.62 | 0.85 | 0.50 | 0.78 | 0.81 |
|  | Correlation | **0.90** | **0.77** | **0.93** | 0.90 | 0.89 | 0.91 |
|  | Water ions | 0.88 | 0.76 | 0.90 | **0.90** | **0.95** | **0.93** |
| RUV | US | 0.94 | 0.57 | 0.89 | 0.90 | 0.90 | 0.95 |
|  | Correlation | 0.94 | 0.61 | 0.93 | 1.00 | 0.90 | 0.93 |
|  | Water ions | 0.93 | 0.73 | 0.92 | 0.50 | 0.78 | 0.74 |
| RUV random | US | 0.96 | 0.68 | 0.92 | 0.90 | 0.95 | 0.95 |
|  | Correlation | 0.96 | 0.69 | 0.90 | 0.80 | 0.90 | 0.92 |
|  | Water ions | **0.91** | **0.73** | **0.93** | **1.00** | **0.95** | **0.96** |

**Supplementary Figure 1:**
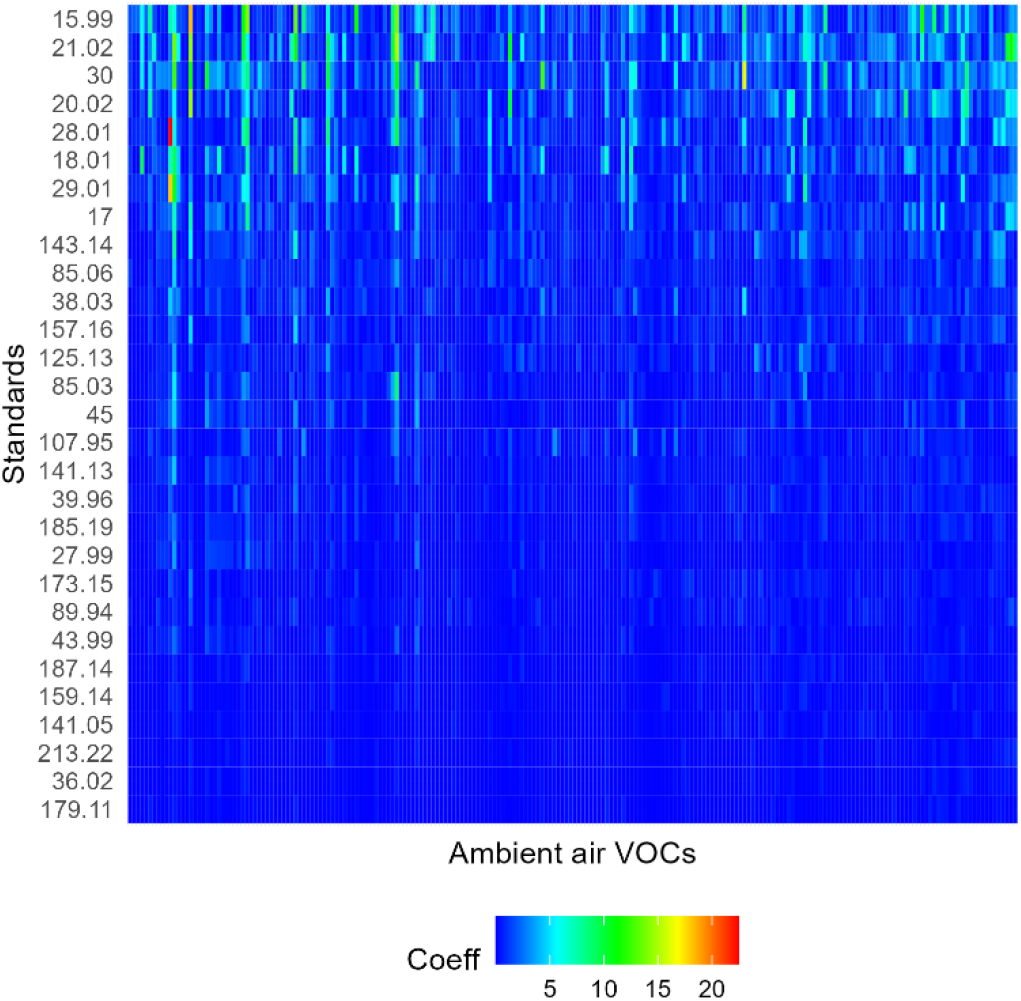
Internal standards selected for the NOMIS method in the emergency department cohort. The plot represents the estimated coefficient for each recording. IS are ordered by coefficient value, *i*.*e*. by importance in normalisation.

